# Investigating the Impact of Habitual Sleep Quality on Episodic Memory Performance: An EEG-Based Representational Similarity Analysis

**DOI:** 10.1101/2024.11.08.622661

**Authors:** Masoud Seraji, Soroush Mirjalili, Audrey Duarte, Vince D Calhoun

## Abstract

Sleep is crucial for episodic memory consolidation, yet the impact of habitual sleep quality on memory performance remains underexplored. This study investigates the relationship between sleep quality and episodic memory retrieval using EEG-based representational similarity analysis (RSA). Thirty-six participants wore wrist accelerometers for one week to capture habitual sleep patterns, including total sleep time and restlessness. Memory performance was assessed through a paired associate learning task, with EEG data recorded during encoding and retrieval phases. RSA was applied to EEG oscillatory power across time-frequency windows to examine the neural similarity between encoding and retrieval. The results showed both positive and negative correlations between sleep metric and memory performance, with sleep restlessness being linked to both increases and decreases in neural similarity across specific clusters. These findings emphasize the important role of sleep quality in shaping the neural processes underlying episodic memory retrieval, indicating a strong connection between sleep patterns and memory function.

## 1 INTRODUCTION

Sleep plays a critical role in consolidating episodic memory [1]. Experimental studies manipulating sleep conditions—such as comparing morning versus evening retrieval, sleep deprivation, or the presence of intervening naps—have shown that episodic memory (i.e., memory of past experiences) is influenced by sleep in both younger and older adults [2]. Additionally, polysomnography research has identified EEG sleep patterns associated with memory consolidation across different age groups [3]. However, these studies are typically conducted in controlled settings and often do not capture natural sleep patterns in a home environment or extend beyond a single night of monitoring.

A widely used method for evaluating habitual sleep is actigraphy, which provides various sleep parameters, including total sleep time, sleep efficiency, and wake after sleep onset. Neurobiological models of memory suggest that successful retrieval of episodic memories relies on the reactivation of neural activity patterns and processes involved during the initial encoding phase [4]. This concept has been supported by neuroimaging research in young adults using encoding-retrieval similarity (ERS) analyses, which demonstrate that reactivation of encoding-related neural patterns reflects task-specific and category-level information crucial for memory accuracy [5]. Additionally, neuroimaging evidence shows that reactivation of event-specific neural patterns, beyond just category or task reactivation, plays a vital role in successful recollection [6], [7]. Although ERS has been linked to episodic memory performance in young adults [6], the role of habitual sleep quality in this relationship remains relatively underexplored. In this study, we collected habitual sleep data over one week using wrist-worn accelerometers, allowing us to capture both average sleep quality and night-to-night variability. This enabled us to investigate how these sleep measures influence episodic memory performance. To assess neural similarity between encoding and retrieval, we applied representational similarity analysis (RSA) to time-frequency EEG data obtained during a paired associate learning task. Specifically, we evaluated event-specific, oscillatory similarities across various frequencies between paired associates during encoding and retrieval across the adult lifespan. Given the established link between sleep quality and memory performance in both young and older adults [3], we hypothesize that individual differences in both memory performance and ERS will be associated with differences in sleep quality. It is important to note that we collected data from both young and older participants; however, the current results are preliminary, and we have not yet analyzed the effect of age in this study.

## 2. METHODS

### 2.1. Participants

The participant sample consisted of 36 right-handed adults (22 females, 12 males, 1 other, 1 transgender-woman), ranging in age from 18 to 74 (young group:18-36, old group: 56-74) years (M = 38, SD = 20). All participants self-reported being native English speakers, right-handed, and having normal or corrected-to-normal vision. None reported any uncontrolled psychiatric, neurological, or sleep disorders, nor vascular disease. Out of the 36 participants, sleep data was missing for three, resulting in a final sample of 33 participants included in the analysis.

### 2.2. Procedure

The experimental design spans seven days, involving lab visits and at-home data collection. On Day 0, participants visit the lab for neuropsychological assessments and the setup of accelerometers (Actiwatch 2, Philips) to track habitual sleep patterns. Over Days 1-2, participants were at home, where nightly sleep data was recorded. On Day 3, they return to the lab for memory encoding and immediate memory retrieval tasks. Days 4-6 involve continued at-home monitoring, followed by a final lab visit on Day 7 for a delayed memory retrieval task (Fig. 1). In the memory task, during the encoding phase, participants were shown images of objects within scenes for 4 seconds, followed by a 350-750ms fixation screen. They were asked whether the object fits within the scene, with possible responses being “Yes” (1), “No” (2), or “Somewhat” (3). In the retrieval phase, participants viewed images and were asked to classify them as either “Same Old” (when both the object and background were unchanged), “Different Old” (when the object was the same, but the background had changed), or “New” (when the object was new) (Fig. 1).

**Fig. 1.**
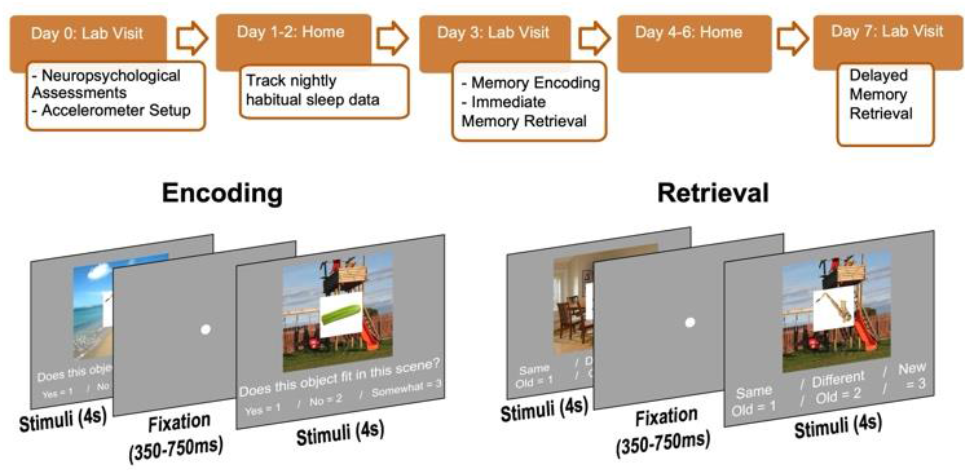
The experimental design consisted of lab visits and at-home sleep tracking using accelerometers. Participants completed memory encoding tasks by judging if an object fit within a scene. During retrieval, they identified whether the object was “Same Old” (both object and background were unchanged), “Different Old” (the object was the same, but the background had changed), or “New” (the object was new). The study spanned 7 days, with habitual sleep data collected between tasks.

### 2.3. EEG acquisition and preprocessing

Continuous EEG data were recorded from 31 scalp electrodes using the Brain Vision ActiCAP system. The electrodes were positioned according to the extended 10–20 system [8], with placements at Fp1, Fz, F3, F7, FT9, FC5, FC1, C3, T7, TP9, CP5, CP1, Pz, P3, P7, O1, Oz, O2, P4, P8, TP10, CP6, CP2, C4, T8, FT10, FC6, FC2, F4, F8, and Fp2. Additionally, two electrodes at the lateral canthi of the right and left eyes captured horizontal electrooculogram (HEOG), while two electrodes placed above and below the right eye recorded vertical electrooculogram (VEOG). The EEG data were sampled at 500 Hz without the application of any high or low pass filters.

EEG data was analyzed offline using MATLAB with the EEGLAB [9] and FIELDTRIP [10] toolboxes. The data underwent baseline correction from 400 to 200 ms prior to stimulus onset, with the sampling rate reduced from 500 Hz to 250 Hz. It was re-referenced to the average of the left and right mastoid electrodes, bandpass-filtered between 0.05 and 80 Hz, and 60 Hz line noise was also removed. Initial artifact removal involved visual inspection to discard trials with muscle, electrode, or sweat artifacts. An independent component analysis (ICA) was then performed to detect and remove ocular artifacts (e.g., blinks, eye movements). Epochs with extreme voltages over 150 microvolts were automatically excluded, with further manual inspection for any remaining artifacts. Finally, time-frequency analysis was conducted using Morlet wavelets [9].

In the wavelet analysis, 38 linearly spaced frequencies between 3 and 40 Hz were used [11], and the data were subsequently down-sampled from 250 Hz to 50 Hz. During the wavelet transformation, each epoch was reduced to the time range of interest (0–2,400 ms post-stimulus onset). As a result, the data dimensions were transformed to n(trials) × 31(electrodes) × 38(frequencies), with power values calculated for 120 time bins of 20 ms each (corresponding to the 50 Hz sampling rate over 2.4 seconds). We grouped the electrodes into four distinct, non-overlapping regions (as illustrated in Fig. 2) and averaged the signals from all electrodes within each region. The wavelet transforms were segmented into 22 time windows of 300 ms each, with consecutive windows overlapping by 100 ms. A detailed summary of these procedures is provided in Fig. 2.

**Fig. 2.**
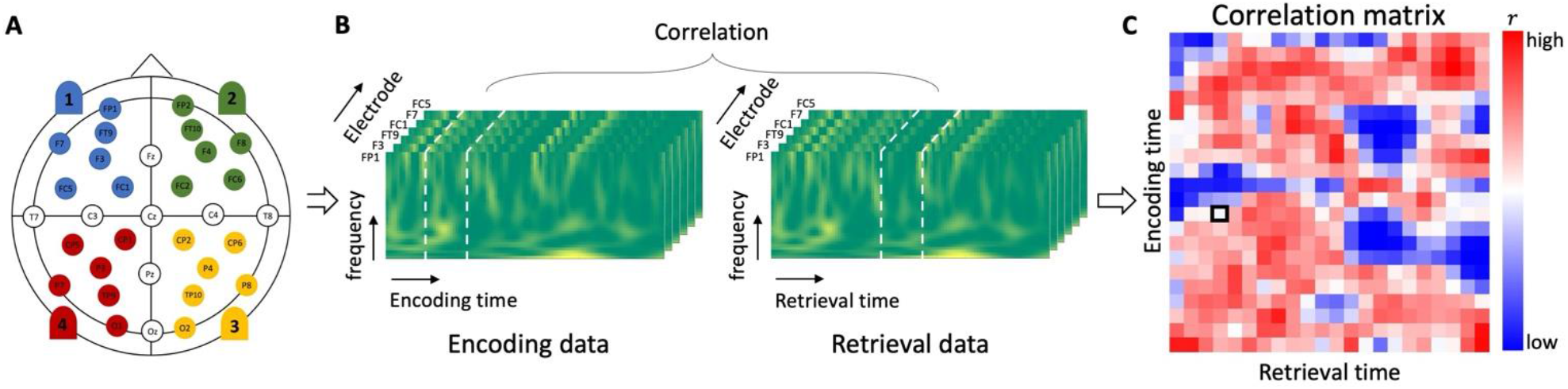
The methodology for the representational similarity analysis (RSA) consisted of three main steps: (A) dividing the brain into four regions (left frontal, right frontal, left posterior, right posterior), and (B) averaging EEG power within each 300 ms time window for each frequency and electrode. These power values were log-transformed and averaged across electrodes within each brain region to generate a vector of 38 log-transformed power values (spanning 3–40 Hz), known as representational patterns. For each brain region, these vectors were further averaged to create a single representative vector for the region. In the figure, the time windows from 300–600 ms during encoding (left pattern) and 800–1,100 ms during retrieval (right pattern) for the left frontal region are highlighted with dashed white lines. (C) The representational patterns from these time windows are correlated using Pearson correlation, with the correlation coefficient displayed as a black square on the matrix. This process is repeated for all time windows during encoding and retrieval, resulting in a full correlation matrix.

### 2.4. RSA Analysis

We used RSA in this study to examine how EEG signals correspond to memory performance during a memory encoding and retrieval task. Participants first encoded visual scenes by determining whether specific objects fit within scene (encoding phase). Later, during retrieval, they were asked to identify whether objects in new/old scenes matched what they had seen during encoding (retrieval phase). EEG oscillatory power from 3 to 40 Hz was measured during both encoding and retrieval, and these power values were averaged within 300 ms time windows for each electrode. The values were then log-transformed and averaged across electrodes in specific brain regions (e.g., left frontal). The key analysis focused on two memory conditions: *hits* and *misses*. A *hit* was defined as correctly identifying a previously attended scene as a match during retrieval, while a *miss* was incorrectly identifying a matched attended scene as a mismatch.

To assess the neural similarity between encoding and retrieval, we computed Pearson correlations between the EEG power vectors in the 300 ms time windows across the encoding and retrieval phases, this approach has been used in prior EEG studies [12], [13], [14]. We then compared within-event similarity (retrieval event and the associated matching encoding event) and between-event similarity (retrieval event compared with all encoding events from the same category). This was done for hits and misses to examine how neural patterns differentiate between successful and unsuccessful memory retrievals. In the next step of the analysis, we subtracted the between-event similarity matrices from the within-event similarity matrices for each trial. We then averaged these event-specific, time–time similarity matrices across trials of the same type for each participant, generating an overall average time–time similarity matrix for each electrode region and trial type. To isolate memory performance differences, we subtracted the average miss similarity from the average hit similarity. Specifically, we calculated the difference between within-event and between-event similarities for hits and misses: (within-hit − between-hit) − (within-miss − between-miss).

### 2.5. Actigraphy data

We extracted key sleep metrics, including total sleep time (TST), sleep efficiency, fragmentation, sleep onset latency, wake after sleep onset (WASO), and the number of wake bouts, for analysis. TST represents the total time spent asleep during a sleep period, while sleep efficiency is the percentage of time spent asleep while in bed. Sleep fragmentation reflects restlessness during sleep, onset latency is the time it takes to fall asleep, WASO captures the total time awake after falling asleep, and wake bouts count the number of awakenings during sleep. We calculated the mean and variance of six sleep metrics over seven days and performed two separate principal component analysis (PCA). These six variables were grouped into two primary components: sleep duration (including total sleep time, sleep efficiency, and inverse sleep onset latency) and restless sleep (including fragmentation, WASO, and wake bouts).

### 2.5. Statistical analysis

To identify significant clusters, correlations between sleep metrics and event-specific encoding-retrieval similarity were calculated for each encoding period to explore the relationship between sleep variability and neural reactivation patterns. Time windows with correlation values exceeding a predetermined threshold (p ≤ 0.05) were selected and grouped into contiguous clusters based on temporal proximity. For example, if the ([200–500 ms] encoding, [500–800 ms] retrieval) and ([300–600 ms] encoding, [500– 800 ms] retrieval) windows both showed significant correlations, they would be combined into a single cluster covering ([200–600 ms] encoding, [500–800 ms] retrieval). It is important to note that this study utilized delayed retrieval, offering further insights into how neural activity patterns over time contribute to long-term memory recall. The significance of these newly combined temporal clusters was assessed by permuting the correlation values 10,000 times, creating a null probability distribution for the cluster-based statistics [15].

## 3. RESULTS

After performing PCA, the analysis reduced correlated variables into uncorrelated components that capture the data’s variance, we identified two primary sleep metrics: sleep duration and restlessness, each defined by its mean and variance, resulting in four total metrics. We then computed the correlation between d-prime [16], a measure of memory performance, and each of the sleep metrics. For further analysis, we selected mean restlessness as the sleep metric most strongly correlated with d-prime.

As described in the methods section, we calculated correlations between sleep metrics and event-specific ERS. The clusters where mean restlessness shows significant correlation are displayed in Fig. 3. For instance, in the left frontal region, we identified two clusters that exhibited negative correlations with mean restlessness. The first cluster showed significant negative correlation during an encoding window of 700–1300 ms and a retrieval window of 0–600 ms. The second cluster, spanning the 700–1100 ms encoding window and the 1300–1800 ms retrieval window, also exhibited a negative correlation with mean restlessness. Beyond the left frontal region, other significant clusters were observed in the right frontal, left posterior, and right posterior regions, with positive correlations (Fig. 3).

**Fig. 3.**
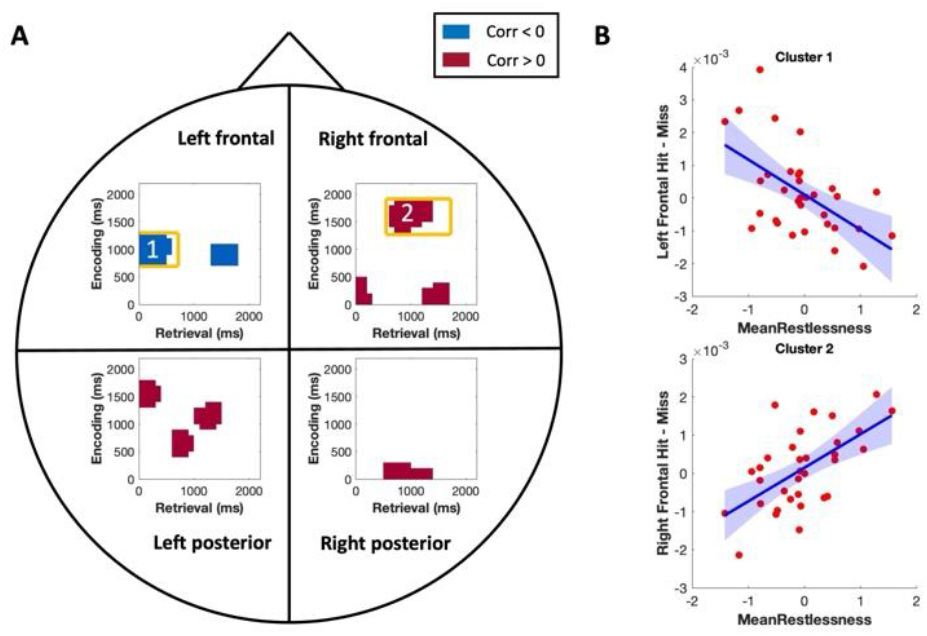
ERS time-time clusters for object-scene pairs displaying correlations with sleep. (A) Each figure highlights the time intervals showing significant correlations between mean sleep restlessness and event-specific ERS. Encoding intervals on the y-axis and retrieval intervals on the x-axis (in m-seconds). Clusters with positive correlations are shown in red, and those with negative correlations are shown in blue. The clusters were corrected using the cluster-based permutation method. (B) The relationship between event-specific ERS and mean restlessness (with corresponding 95% confidence intervals) is illustrated for two of the clusters shown in part (A). These two clusters are provided as examples, one demonstrating a positive correlation and the other a negative correlation, with similar patterns observed across the remaining clusters.

## 4 DISCUSSION AND CONCULSION

Successful episodic memory retrieval is thought to rely on the reactivation of neural activity patterns and associated cognitive processes that were present during the original encoding of the event [4]. In this study, we used EEG to investigate episodic neural reinstatement effects that are sensitive to image-pair-specific activity patterns. We found that neural activity during encoding and retrieval was correlated within similar time intervals. Interestingly, we also observed significant *asymmetries* in ERS effects, where encoding activity was correlated with retrieval activity occurring either earlier or later in time. Several factors could explain these asymmetrical ERS patterns. One possibility is that encoding-related activity, and its associated cognitive processes may persist longer during encoding than during retrieval, leading to later encoding intervals correlating with earlier retrieval intervals. This explanation aligns with findings suggesting that episodic reinstatement occurs on a temporally compressed timescale relative to the original encoding process [17]. Another, non-mutually exclusive explanation is that some cognitive processes occurring earlier during encoding, such as perceptual encoding, may be reinstated later during retrieval, particularly during efforts to recall previously learned word image or to reject rearranged ones [18]. Moreover, these symmetrical and asymmetrical reinstatement effects appeared to be linked to variations in sleep quality. Specifically, sleep restlessness in sleep patterns may influence the efficiency and timing of these neural reactivation processes, potentially affecting how effectively memory traces are accessed and reinstated during retrieval. This extends previous findings by highlighting the role of sleep in modulating not just overall memory performance, but also the temporal dynamics of neural reinstatement that underpin successful episodic retrieval. Furthermore, in clusters showing a negative correlation, higher restlessness (indicating poor sleep) is associated with worse memory performance [19]. On the other hand, in clusters with a positive correlation between ERS, a potential explanation for these results could lie in the distinct roles that familiarity and recollection play in the associative recognition of object-scene pairs [13]. This aligns with previous studies that found both positive and negative correlations between sleep metrics and ERS [13]. Future research should focus on exploring how these sleep-memory relationships differ across age groups, particularly whether the connection between sleep quality and memory is stronger or weaker in older adults compared to younger individuals.

In summary, the findings reinforce the link between sleep quality and episodic memory retrieval, highlighting how sleep affects both the occurrence and timing of neural reactivation. These insights could inform interventions to enhance memory, particularly for individuals with sleep disturbances.

## 5 COMPLIANCE WITH ETHICAL STANDARDS

All participants signed consent forms approved by the University of Texas at Austin Institutional Review Board.

## 6. ACKNOWLEDGMENTS

This study was funded by NSF 2112455 and NSF 2152492.

## Notes

### Competing Interest Statement

The authors have declared no competing interest.

## REFERENCES

[1] B. Rasch and J. Born, “About sleep’s role in memory,” Physiol Rev, vol. 93, no. 2, pp. 681–766, 2013, doi: 10.1152/PHYSREV.00032.2012.

[2] J. M. Wilson, P. J. Marin, M. R. Rhea, S. M. C. Wilson, J. P. Loenneke, and J. C. Anderson, “Concurrent training: a meta-analysis examining interference of aerobic and resistance exercises,” J Strength Cond Res, vol. 26, no. 8, pp. 2293–2307, Aug. 2012, doi: 10.1519/JSC.0B013E31823A3E2D.

[3] B. A. Mander, J. R. Winer, and M. P. Walker, “Sleep and Human Aging,” Neuron, vol. 94, no. 1, p. 19, Apr. 2017, doi: 10.1016/J.NEURON.2017.02.004.

[4] K. A. Norman and R. C. O’Reilly, “Modeling Hippocampal and Neocortical Contributions to Recognition Memory: A Complementary-Learning-Systems Approach,” Psychol Rev, vol. 110, no. 4, pp. 611–646, Oct. 2003, doi: 10.1037/0033-295X.110.4.611.

[5] E. M. Gordon, T. O. Laumann, B. Adeyemo, J. F. Huckins, W. M. Kelley, and S. E. Petersen, “Generation and Evaluation of a Cortical Area Parcellation from Resting-State Correlations,” Cereb Cortex, vol. 26, no. 1, pp. 288–303, Jan. 2016, doi: 10.1093/CERCOR/BHU239.

[6] E. A. Wing, M. Ritchey, and R. Cabeza, “Reinstatement of individual past events revealed by the similarity of distributed activation patterns during encoding and retrieval,” J Cogn Neurosci, vol. 27, no. 4, pp. 679–691, Apr. 2015, doi: 10.1162/JOCN_A_00740.

[7] M. Silva, C. Baldassano, and L. Fuentemilla, “Rapid Memory Reactivation at Movie Event Boundaries Promotes Episodic Encoding,” Journal of Neuroscience, vol. 39, no. 43, pp. 8538–8548, Oct. 2019, doi: 10.1523/JNEUROSCI.0360-19.2019.

[8] M. R. Nuwer et al., “IFCN standards for digital recording of clinical EEG,” Electroencephalogr Clin Neurophysiol, vol. 106, no. 3, pp. 259–261, Mar. 1998, doi: 10.1016/S0013-4694(97)00106-5.

[9] A. Delorme and S. Makeig, “EEGLAB: An open source toolbox for analysis of single-trial EEG dynamics including independent component analysis,” J Neurosci Methods, vol. 134, no. 1, pp. 9–21, Mar. 2004, doi: 10.1016/j.jneumeth.2003.10.009.

[10] R. Oostenveld, P. Fries, E. Maris, and J. M. Schoffelen, “FieldTrip: Open source software for advanced analysis of MEG, EEG, and invasive electrophysiological data,” Comput Intell Neurosci, vol. 2011, 2011, doi: 10.1155/2011/156869.

[11] S. Hanslmayr, B. P. Staresina, and H. Bowman, “Oscillations and Episodic Memory: Addressing the Synchronization/Desynchronization Conundrum,” Trends Neurosci, vol. 39, no. 1, pp. 16–25, Jan. 2016, doi: 10.1016/J.TINS.2015.11.004/ASSET/D27E9EB7-A324-4EA3-9506-0FEA1B136299/MAIN.ASSETS/GR2.JPG.

[12] V. R. Sommer, Y. Fandakova, T. H. Grandy, Y. L. Shing, M. Werkle-Bergner, and M. C. Sander, “Neural Pattern Similarity Differentially Relates to Memory Performance in Younger and Older Adults,” Journal of Neuroscience, vol. 39, no. 41, pp. 8089–8099, Oct. 2019, doi: 10.1523/JNEUROSCI.0197-19.2019.

[13] E. Hokett, S. Mirjalili, and A. Duarte, “Greater sleep variance related to decrements in memory performance and event-specific neural similarity: a racially/ethnically diverse lifespan sample,” Neurobiol Aging, vol. 117, pp. 33–43, Sep. 2022, doi: 10.1016/J.NEUROBIOLAGING.2022.04.015.

[14] S. A. Justus, S. Mirjalili, P. S. Powell, and A. Duarte, “Neural reinstatement of context memory in adults with autism spectrum disorder,” Cerebral Cortex, vol. 33, no. 13, pp. 8546–8556, Jun. 2023, doi: 10.1093/CERCOR/BHAD139.

[15] E. Maris and R. Oostenveld, “Nonparametric statistical testing of EEG- and MEG-data,” J Neurosci Methods, vol. 164, no. 1, pp. 177–190, Aug. 2007, doi: 10.1016/J.JNEUMETH.2007.03.024.

[16] B. C. Haatveit, K. Sundet, K. Hugdahl, T. Ueland, I. Melle, and O. A. Andreassen, “The validity of d prime as a working memory index: results from the ‘Bergen n-back task,’” J Clin Exp Neuropsychol, vol. 32, no. 8, pp. 871– 880, Oct. 2010, doi: 10.1080/13803391003596421.

[17] K. Yaffe, C. M. Falvey, and T. Hoang, “Connections between sleep and cognition in older adults,” Lancet Neurol, vol. 13, no. 10, pp. 1017–1028, Oct. 2014, doi: 10.1016/S1474-4422(14)70172-3/ASSET/F8D83E3F-925E-4333-9968-9D062170AC04/MAIN.ASSETS/GR2.GIF.

[18] A. Jafarpour, L. Fuentemilla, A. J. Horner, W. Penny, and E. Duzel, “Replay of Very Early Encoding Representations during Recollection,” Journal of Neuroscience, vol. 34, no. 1, pp. 242–248, Jan. 2014, doi: 10.1523/JNEUROSCI.1865-13.2014.

[19] B. A. Mander, J. R. Winer, W. J. Jagust, and M. P. Walker, “Sleep: A novel mechanistic pathway, biomarker, and treatment target in the pathology of Alzheimer’s disease?,” Trends Neurosci, vol. 39, no. 8, p. 552, Aug. 2016, doi: 10.1016/J.TINS.2016.05.002.

